# Congruency effects can compensate for deficits of healthy older adults in crossmodal integration

**DOI:** 10.1101/673491

**Authors:** Focko L. Higgen, Charlotte Heine, Lutz Krawinkel, Florian Göschl, Andreas K. Engel, Friedhelm C. Hummel, Gui Xue, Christian Gerloff

## Abstract

One of the pivotal challenges of aging is to maintain independence in the activities of daily life. In order to adapt to changes in the environment, it is crucial to continuously process and accurately combine simultaneous input from different sensory systems, i.e., crossmodal integration.

With aging, performance decreases in multiple cognitive domains. The processing of sensory stimuli constitutes one of the key features of this deterioration. Age-related sensory impairments affect all modalities, substantiated by decreased acuity in visual, auditory or tactile detection tasks.

However, whether this decline of sensory processing leads to impairments in crossmodal integration remains an unresolved question. While some researchers propose that crossmodal integration degrades with age, others suggest that it is conserved or even gains compensatory importance.

To address this question, we compared behavioral performance of older and young participants in a well-established crossmodal matching task, requiring the evaluation of congruency in simultaneously presented visual and tactile patterns. Older participants performed significantly worse than young controls in the crossmodal task when being stimulated at their individual unimodal visual and tactile perception thresholds. Performance increased with adjustment of stimulus intensities. This improvement was driven by better detection of congruent stimulus pairs (p<0.01), while detection of incongruent pairs was not significantly enhanced (p=0.12).

These results indicate that age-related impairments lead to poor performance in complex crossmodal scenarios and demanding cognitive tasks. Performance is enhanced when inputs to the visual and tactile systems are congruent. Congruency effects might therefore be used to develop strategies for cognitive training and neurological rehabilitation.

## Introduction

As the percentage of older people in the population increases, aging related declines gain more and more significance. An important agenda therefore is to identify means for supporting older adults to maintain sound minds and independent living.

In order to behave adequately in our natural environment, it is crucial to continuously process simultaneous input from different sensory systems and integrate this information to meaningful percepts (1). This crossmodal integration complements unimodal sensory perception and allows for basing decisions and behavior on a broader range of sensory cues (2).

Age-related decline affects processes of crossmodal integration in several ways. The processing of sensory stimuli constitutes one of the key features of this deterioration. Age-related sensory impairments affect all modalities, as mirrored in decreased acuity in visual, auditory or tactile detection tasks (3–8). Apart from a decline of peripheral sensory organs, older adults show deficits in attention, working memory, episodic memory, decision making and subsequent actions (9–11). These cognitive domains are highly reliant on the crossmodal perception of information and vice versa.

However, the relevance of crossmodal integration in older adults is still under debate (see for example 12–15). While some authors report that the neurocomputational integration of multiple sensory stimuli degrades with age (e.g. 16–18), others suggest that crossmodal integration is conserved or even gains compensatory importance in older adults (e.g. 19–22).

Classically, the decline in sensory organs and higher cognitive domains was thought to prevent older adults from taking advantage of crossmodal information, by restricting effective multisensory integration processes and limiting the cognitive resources needed (e.g. 18).

More recent evidence, however, points to enhanced crossmodal integration in older adults. Different mechanisms have been discussed as possible reasons for this enhancement. Freiherr and colleagues (15) argue that crossmodal information can compensate for the age-related impairments of peripheral sensory organs. According to the principle of inverse effectiveness (23,24) crossmodal integration can lead to maximal behavioral enhancement during perception of weak or ambiguous stimuli, due to appropriate weighting of the sensory evidence. Consequently, the integration of multiple sensory inputs could ameliorate unimodal decline in older adults’ sensory systems and improve the overall outcome of perceptual processing (25).

Age-related alterations in central neurocomputational processes have been discussed to influence crossmodal integration. General cognitive slowing in older adults, demonstrated in several tasks (26–28), has been suggested to lead to more susceptibility to crossmodal integration in older adults by extending the temporal window for possible crossmodal interactions (29,30). Furthermore, it has been proposed that gains in performance in scenarios with crossmodal stimulation (31,32) might relate to increases in baseline crossmodal interactions in older adults due to neural noise (33,34).

To further characterize the relevance of crossmodal integration in older adults, we investigated group differences between healthy older and young participants in a well-established visuo-tactile matching task (35–38). In this task, participants have to evaluate congruency in simultaneously presented visual and tactile dot patterns. Most studies that found a behavioral benefit of older adults in crossmodal tasks have focused on visual-auditory integration (20–22). Data on other sensory modalities are sparse. Our study focuses on visuo-tactile integration, as there is evidence suggesting that tasks involving interaction of visual and somatosensory stimuli profit strongly from crossmodal interactions effects (39,40). In everyday life, the tactile modality interacts mainly with vision to recognize our surroundings and facilitate orientation. The latter represents a basic ability needed for interacting with the environment and to preserve independence (41,42).

To be able to compare participants’ performance and subjective task difficulty across both groups and modalities, we determined individual unimodal perception thresholds prior to the crossmodal study (43,44).

Our first hypothesis was that unimodal perception thresholds for visual and tactile pattern recognition should be higher in the older group compared to young, due to multiple age-related sensory impairments (45). Second, we hypothesized that older participants would show greater enhancement of performance compared to young in a crossmodal integration task involving stimuli presented at the individual unimodal perceptual thresholds – in accordance with the more recent notion in the field.

## Materials and Methods

### Participants

37 older and 22 young volunteers were screened for the study. Six older volunteers did not meet the inclusion criteria during initial assessment. One older and two young participants dropped out because of personal reasons or technical problems. Ten older participants (five female, mean (M) = 74.1 years, standard deviation (SD) = 3.90 years) did not meet the predefined accuracy criterion in a training session prior to the threshold estimation (described in detail in ‘Experimental procedure’) and were no longer considered in the analysis. Thus, the final sample consisted of 20 young (11 female, M = 24.05 years, SD = 2.50) and 20 older (11 female, M = 72.14 years, SD = 4.48) volunteers. All participants were right-handed according to the Edinburgh handedness inventory (Oldfield, 1971), had normal or corrected to normal vision, no history or symptoms of neuro-psychiatric disorders (MMSE ≥ 28, DemTect ≥ 13) and no history of centrally acting drug intake. All participants received monetary compensation for participation in the study.

### Assessment

Prior to inclusion, each participant underwent an assessment procedure. Assessment consisted of a neurological examination, the Mini-Mental State Examination (MMSE; 46) and the DemTect (47) to rule out symptoms of neuro-psychiatric disorders. Furthermore, a 2-point-discrimination test (cut-off > 3mm; 48,49) and a test of the mechanical detection threshold (cut-off > 0.75mN; MDT, v. Frey Filaments, OptiHair2-Set, Marstock Nervtest, Germany; 50,51) were conducted to ensure intact peripheral somatosensation.

### Setup and stimuli

The experiment was conducted in preparation for a magnetoencephalography (MEG) study and the setup was designed to match conditions in the MEG laboratory. The experiment took place in a light-attenuated chamber. We chose experimental procedure, stimulus configuration and stimulation parameters based on pilot data showing accuracy of tactile pattern recognition to be very different between older and young participants.

We used an adapted version of a well-established experimental paradigm, the visuo-tactile matching task (35–38). Participants are instructed to compare tactile patterns presented to the right index fingertip and visual patterns presented on a computer screen. For tactile stimulation, the participants’ right hand was resting on a custom-made board containing a Braille stimulator (QuaeroSys Medical Devices, Schotten, Germany, see Figure 1). The Braille stimulator consists of eight pins arranged in a four-by-two matrix, each 1mm in diameter with a spacing of 2.5mm. Each pin is controllable independently. Pins can be elevated for any period of time to form different patterns. At the end of each pattern presentation, all pins return to baseline. The stimuli consisted of four geometric patterns, each of them formed by four elevated pins (Figure 1). Participants passively perceived the elevated pins without active exploration. A 15-inch screen at 60Hz with a resolution of 1024×768 pixels, positioned 65cm in front of the participants, served for presentation of the visual stimuli. The design of the visual patterns was analogous to the tactile patterns. Visual patterns had a size of 43×104 pixels. They were presented left of a central fixation point on a noisy background (Perlin noise; Figure 1D).

**Fig. 1.**
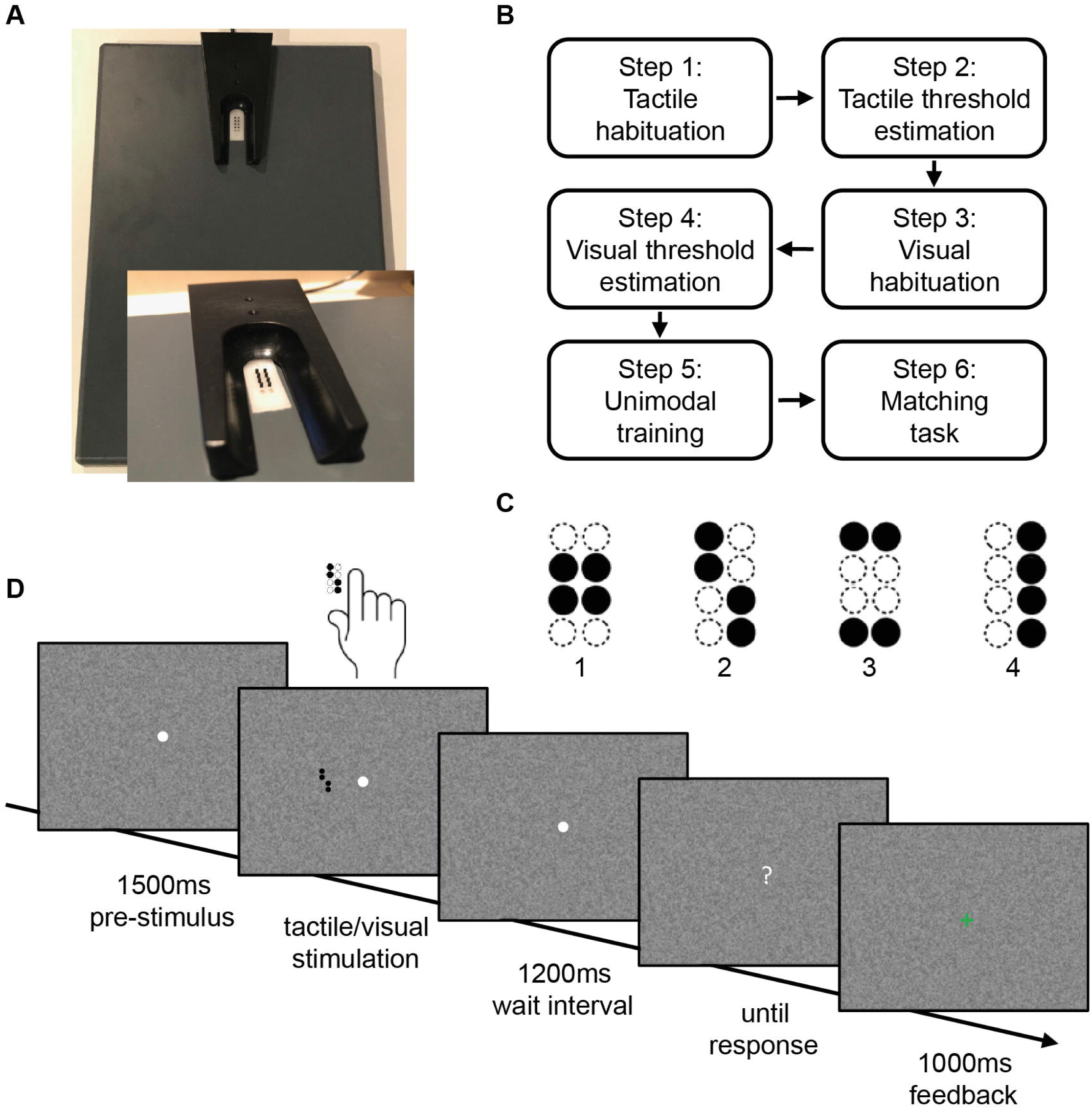
Stimulus design and experimental procedure. **A**: Braille stimulator. For tactile stimulation, the participants’ right hand was resting on a custom-made board containing the Braille stimulator (QuaeroSys Medical Devices, Schotten, Germany), with the fingertip of the right index finger placed above the stimulating unit. **B**: Sequence of tasks in the experiment, **C**: Stimuli consisted of four different patterns, **D**: After a pre-stimulus interval of 1500ms, tactile and/or visual patterns were presented for 500ms depending on the current step of the experiment. After a wait interval of 1200ms, a question mark appeared on the screen and participants gave the response via button press. Depending on the current step of the experiment, visual feedback was given (1000ms).

Depending on the task, the amplitude of pin elevation and the grey intensity of visual patterns were adjusted, while the duration of the pattern presentation was always kept constant at 500ms. The amplitude of the pin elevation can be controlled in 4095 discrete steps, with a maximum amplitude of 1.5mm. Maximal grey intensity equaled black patterns with RGB: 0-0-0.

We used Presentation software (Neurobehavioral Systems, version 15.1) to control stimulus presentation and to record participants’ response time (RT) and accuracies.

### Experimental procedure

All participants who met the predefined accuracy criterion in a training session prior to the experiment (at least 75% correct answers in a tactile-to-visual delayed match-to-sample task with easy tactile patterns) performed a series of tasks representing the current experiment (tactile habituation, tactile threshold estimation, visual habituation, visual threshold estimation, unimodal training, matching task; see Figure 1B). At the beginning of each task participants read the task instructions presented on a computer screen. The experiment started with the tactile habituation task.

#### Tactile habituation

The tactile habituation task consisted of a tactile-to-visual delayed match-to-sample task. Four target patterns were introduced as the stimulus set (see Figure 1B, trial sequence), at maximum pin amplitude and with a duration of 500ms. Each trial started with a central white fixation point appearing on a noisy background. This fixation point remained visible throughout each trial. The tactile pattern presentation started 1500ms after appearance of the fixation point with a stimulus chosen pseudo-randomly from the stimulus set. After the tactile presentation and a waiting interval of 1200ms, the central fixation point turned into a question mark and participants indicated which of the four patterns had been presented. Participants responded via button press with the fingers two to five of the left hand. After each trial participants received visual feedback (1000ms) whether their response was correct (green ‘+’) or incorrect (red ‘-’; see Figure 1C). The waiting interval after stimulus offset was integrated to prepare for the following MEG experiment, where it allowed for avoiding motor artefacts in the MEG signal. The background changed after every trial. After a minimum of five training blocks, each consisting of 16 trials, and an accuracy of at least 75% in three of five consecutive blocks, participants could proceed to the next step. If participants did not reach the target accuracy within 15 blocks, they were excluded from further participation.

#### Tactile threshold estimation

Pilot studies indicated that most older adults were able to recognize the target patterns at 500ms stimulus presentation time in the unimodal tactile condition with an accuracy of approximately 80% correct. However, using the same parameters, young performed close to 100%. Equally, visual recognition accuracy was close to 100% in both groups for these parameters. To achieve a comparable performance of around 80% correct answers for both modalities in older and young participants, we conducted an adaptive staircase procedure to detect thresholds for visual and tactile pattern recognition and tailor stimulus intensities for each participant.

Since the slope of the psychometric function was supposed to be very different in older and young participants and we did not have any priors regarding the exact shape, we decided not to use a Bayesian approach (e.g. Quest; 52), but to implement a non-parametric adaptive staircase procedure (53–56). We designed a two-down/one-up fixed-step-size adaptive staircase. With a ratio Δ-down/Δ-up = 0.5488, this staircase should converge around 80.35%. For tactile pattern presentation, adaptation of the height of the braille pins rendered recognition easier or more complicated. Step size was determined after piloting with approximately 0.1mm up, 0.055mm down. The staircase started with the maximum amplitude of 1.5 mm. The staircase stopped after 20 reversals, while proceeding at boundary levels. The last 16 reversals served to calculate thresholds. Participants performed this staircase for both unimodal visual and unimodal tactile stimulation. Trial timing was the same as in the habituation task, except there was no feedback given.

#### Visual habituation

The visual habituation task followed the same procedure as in the tactile condition. Instead of tactile stimulation, patterns were presented visually at maximal contrast (see Figure 1C, target patterns). Again, participants continued to fixate the central point during pattern presentation, so that visual patterns would appear in the left visual hemi-field. Trial timing, block design and accuracy criterion were the same as for the tactile recognition task.

#### Visual threshold estimation

The visual threshold estimation followed the same procedure as in the tactile modality. For visual threshold estimation, adaptation of the grey intensity of the pattern varied the patterns’ contrast against the noisy background. Step size was determined after piloting, with a step up being two intensities, and a step down one on the grey scale ranging from 47 (RGB: 138-138-138) to 101 (RGB: 0-0-0). The staircase started with the maximum contrast (black pattern; RGB: 0-0-0). Pilot data showed that a grey intensity of RGB: 138-138-138, which corresponds to the mean of the grey values of our noisy background, was hardest to detect. Therefore, this contrast was the lower boundary of the staircase. Trial timing and stimulus duration remained the same as in the tactile threshold estimation process.

Following the threshold estimation in tactile and visual modalities, participants performed a short *unimodal training* in both conditions to verify thresholds calculated from the adaptive staircase procedure. The order of modalities was chosen randomly. Trial timing remained the same as in the habituation tasks. To keep performance at a comparable level, thresholds were adjusted if accuracy was below 75% or above 85% over 5 blocks. For the adjustment, the same step sizes as in the adaptive staircase were used.

#### Visuo-tactile matching

After the unimodal threshold estimation, participants conducted the visuo-tactile matching task. In this task, visual and tactile patterns were presented with synchronous onset and offset, and participants had to decide whether the patterns were congruent or incongruent. Participants responded with the left index (‘congruent’) or middle finger (‘incongruent’) via button press on a response box and again visual feedback (a green ‘+’ or a red ‘-’) was given in every trial. Trial timing was the same as in the unimodal recognition task (Figure 1C). Congruent and incongruent stimulus pairs were presented equally often. Participants started the visuo-tactile matching task at the stimulus intensity of the unimodal thresholds and performed a set of five consecutive blocks, consisting of eight trials. If participants did not reach an average accuracy between 75% and 85% correct within these five blocks, stimulus intensities in both modalities were adapted according to the steps of the adaptive staircase procedure.

Visual and tactile stimulus intensities were adjusted evenly. After adjustment of stimulus intensities participants performed another set of five blocks. The experiment ended when participants reached a stable performance between 75-85% correct averaged over a set of five blocks (mean number of sets = 2.25, standard deviation = 0.93).

### Statistical analysis

Statistical analyses were performed using Matlab (Version 8.4.0.150421, MathWorks, Natick, MA, 2014) and RStudio (Version 3.5.4, R Core Team, 2017).

To test for baseline group differences a multivariate analysis of variance (MANOVA) was performed by means of R’s *manova()* command to investigate the relationship between the values for sex, MDT, 2-point-discrimination, MMSE, DemTect as dependent variables and group (young group vs. older group) as the independent variable. As group allocation was defined by participants’ age, age was not included into the model. For post-hoc analysis, two-sampled t-tests were performed and Benjamini-Yekutieli (BY) correction was applied to adjust for multiple comparisons (57). To compare the older and young groups’ performance in the habituation tasks, another MANOVA was performed. Dependent variables were the response accuracies in tactile and visual pattern recognition. For post-hoc analysis, two-sampled t-tests were performed and BY correction was applied to adjust for multiple comparisons. Another MANOVA was performed to compare accuracies and thresholds before and after the unimodal training and between the two groups, whereby accuracies, grey intensities and pin heights were dependent variables and time (before vs. after the training) and group (young group vs. older group) independent variables. Two-sample t-tests and BY correction were used for post-hoc analysis of significant group effects.

As in the course of the visuo-tactile matching task pin height and grey intensity were adjusted evenly according to the steps of the adaptive staircases, changes in stimulus intensities were highly dependent. We therefore did not use MANOVA for further analyses. Two sample t-tests and BY correction were used to compare accuracies and stimulus intensities between the groups in the first and last set of five blocks of the visuo-tactile matching task. In addition, paired t-tests and BY correction were performed to compare accuracies and stimulus intensities within the groups between the first and the last set of the visuo-tactile matching task.

To evaluate detection performance for congruent and incongruent stimulus pairs, paired t-tests were used to compare accuracies within and two-sampled t-tests to compare accuracies between groups. BY correction was used to adjust for multiple comparisons.

## Results

### Baseline data

The group comparison of baseline data obtained in the assessment prior to inclusion (Table 1) showed significant differences between the groups of young and older participants (Pillai’s Trace=0.43, *F*=5.15, *df*=(1,38), *p*<0.01). Post-hoc comparison of the baseline data showed that DemTect scores (*p*<0.001) differed significantly between groups. Importantly, the measurements revealed age-appropriate, not pathological results in the older group.

**Tab. 1.** Baseline data of the groups. Mean values are shown ± standard deviation. Based on a significant main effect of the factor group (young group vs. older group), post-hoc tests were conducted. * indicate significant difference between young and older participants, p-value ≤ 0.01.

| Metrics | Young group (n=20) | Older group (n=20) |
| --- | --- | --- |
| DemTect | 17.8 ($\pm 0.6$)* | 15.9 ($\pm 1.5$)* |
| MMSE | 29.7 ($\pm 0.6$) | 29.6 ($\pm 0.6$) |
| 2-Point (mm) | 2.1 ($\pm 0.2$) | 2.2 ($\pm 0.4$) |
| MDT (mN) | 0.28 ($\pm 0.1$) | 0.57 ($\pm 0.5$) |

### Habituation tasks

The analysis of performance in the habituation tasks revealed significant associations between accuracy and group (Pillai’s Trace=0.41, *F*=12.65, *df*=(1,38), *p*<0.001). Post-hoc comparison showed that the young participants (95.70 ± 5.10%) performed significantly better in the tactile task compared to the older group (82.47 ± 10.44%), (*F*(38)=25.93, *p*<0.001). In the visual task, response accuracies did not differ between the groups (Young group: 98.12 ± 2.94%/Older group 98.12 ± 2.94%; *F*(38)=0, *p*=1).

### Threshold estimation and unimodal training

The results of the threshold estimation are displayed in Figure 2.

**Fig. 2.**
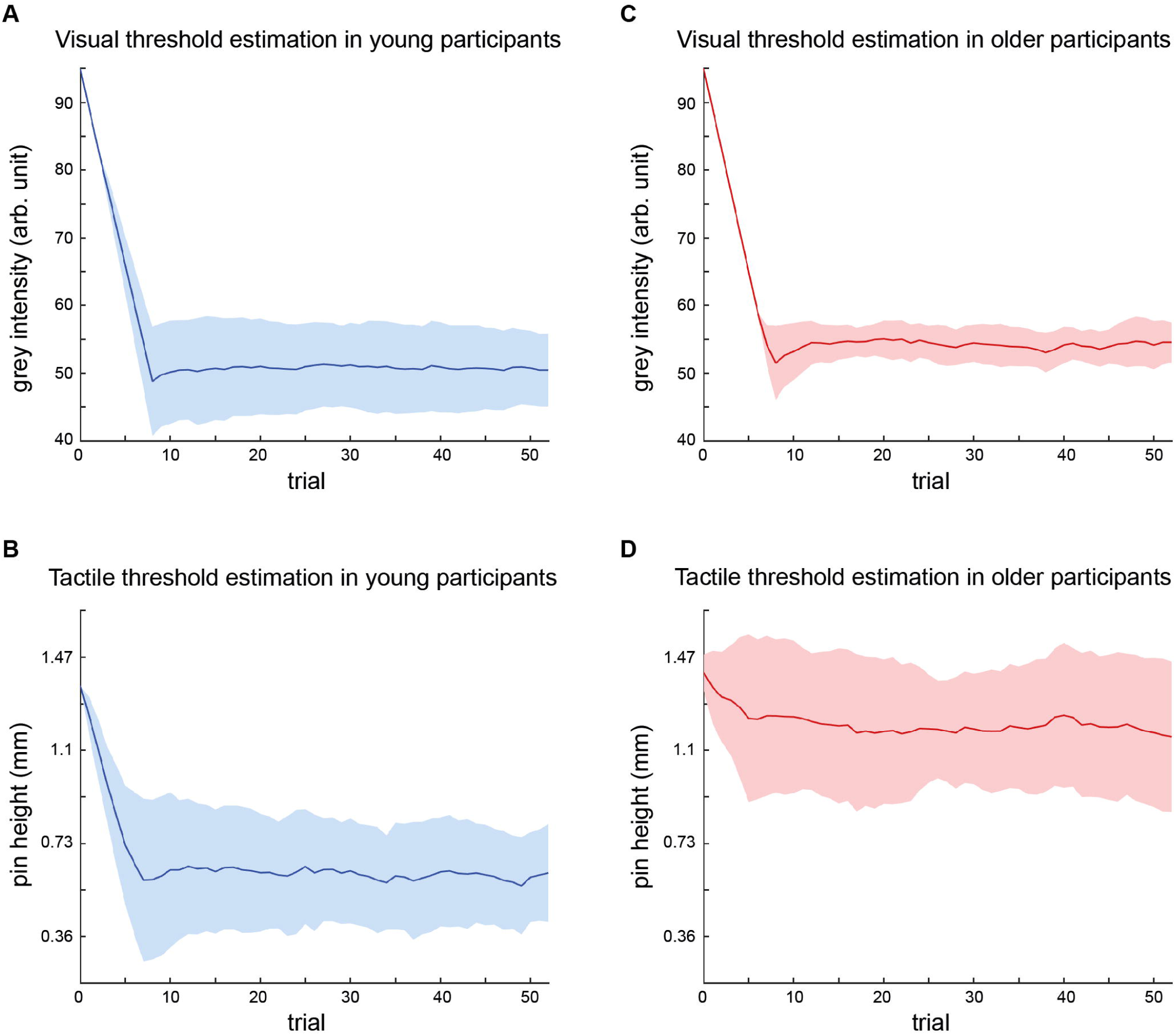
Summary of threshold estimations for visual and tactile stimulus intensities. Graphs depict the mean stimulus intensity (y-axis) per trial (x-axis) during the course of the adaptive staircase over all participants (young group = blue; older group = red) with standard deviations (hatched areas). Number of trials equals trials in shortest threshold estimation procedure, i.e. trials common to all participants. **A**: Visual threshold estimation in young participants, B: Tactile threshold estimation in young participants, **C**: Visual threshold estimation in older participants, **D**: Tactile threshold estimation in older participants.

Visual threshold estimation in the young group resulted in a mean grey intensity of 49.2 ± 1.1. The mean adaptive staircase for tactile threshold estimation in the young group showed a course similar to the visual condition and resulted in a mean threshold, i.e., pin height of 0.60 ± 0.17mm.

Visual threshold estimation in the older group resulted in a mean grey intensity of 53.9 ± 2.5. The mean adaptive staircase for tactile threshold estimation in the older group showed only a small downward trend, indicating that the tactile threshold in the older group was close to maximum stimulus intensity. The mean tactile threshold in the older group was 1.13 ± 0.28mm.

To ensure the validity of the estimated thresholds, the unimodal training was performed. Within the groups, there was no significant change of visual or tactile thresholds in the course of the training (Pillai’s Trace=0.68, *F*=38.51, *df*=(1,77), *p*=0.981), indicating a reliable threshold estimation. Across groups, there was no difference in detection accuracy (tactile *F*(77)=4.29, *p*=0.116, visual *F*(77)=0.72, *p*=0.825), but as expected in pin height (*F*(77)=112.86, *p*<0.001) and grey intensity (*F*(77)=104.77, *p*<0.001).

### Visuo-tactile matching

The mean accuracies, tactile (pin heights) and visual (grey intensities) stimulus intensities of the first and last set of five blocks of the visuo-tactile matching task were compared within and between groups (Table 2).

**Tab. 2.** First and last set of the visuo-tactile matching task. Mean values are shown ± standard deviation for accuracy, grey intensity and pin height in the first and last set of the task, sorted by group. * indicate significant difference between young and older participants, all p-values ≤ 0.001; ^#^ indicate significant difference within the older group, all p-values ≤ 0.01.

|  | Young group (n=20) | Older group (n=20) |
| --- | --- | --- |
| First set of matching task |  |  |
| Accuracy (%) | 78.31 (± 9.09)* | 66.20 (± 9.31)*# |
| Pin height (mm) | 0.58 (± 0.17)* | 1.14 (± 0.28)*# |
| Grey intensity | 49 (± 1.38)* | 53.65 (± 2.70)*# |
| Last set of matching task |  |  |
| Accuracy (%) | 79.50 (± 5.94) | 77.28 (± 6.00)# |
| Pin height (mm) | 0.57 (± 0.17)* | 1.24 (± 0.27)*# |
| Grey intensity | 48.9 (± 1.59)* | 56.2 (± 2.69)*# |

In the first set, older participants performed significantly worse (*t*(37.98)=4.16, *p*<0.01) despite their significantly higher unimodal stimulus intensities (grey intensity: *t*(19.17)=156.69, *p*<0.001; pin height: *t*(30.58)=-7.70, *p*<0.001). To reach a performance of around 80% correct responses in the older group, visual and tactile intensities had to be further increased significantly according to the steps of the adaptive staircase (visual: *t*(19)=-6.71, *p*<0.001, tactile: *t*(19)=-7.75, *p*<0.001). With this adjustment of stimulus intensities task performance was significantly improved (*t*(18)=-4.59, *p*<0.01) and there was no longer a significant difference in accuracy between the young and the older group (*t*(36.86)=1.16, *p*=0.92). Within the young group, there was no difference between the first and the last set in accuracy (*t*(19)=-0.55, *p*=1), grey intensity (*t*(19)=0.81, *p*=1) and pin height (*t*(19)=2.37, *p*=0.12).

### Congruent vs. incongruent stimulus pairs

To further explore the differences in performance in the visuo-tactile matching task, we analyzed detection accuracy for congruent and incongruent stimulus pairs separately.

Both age groups exhibited strong congruency effects with the detection rate for congruent patterns being significantly higher than for incongruent pairs over all matching blocks (older group: congruent 82.12%, incongruent 62.21%, *t*(19)=7.81, *p<*0.001; young group: congruent 87.34%, incongruent 70.33%, *t*(19)=5.51, *p<*0.001). While overall detection accuracy was significantly lower in the older group in the first set of the visuo-tactile matching task (see 3.4), the difference in detection accuracy for congruent vs. incongruent stimulus pairs was the same (18%) for both age groups (older group 75.48% vs. 57.11%, *p*<0.01/young group 87.05% vs. 69.30%, *p*<0.01; no difference between groups in percentage difference, *p*=0.88; see Figure 3).

**Fig. 3.**
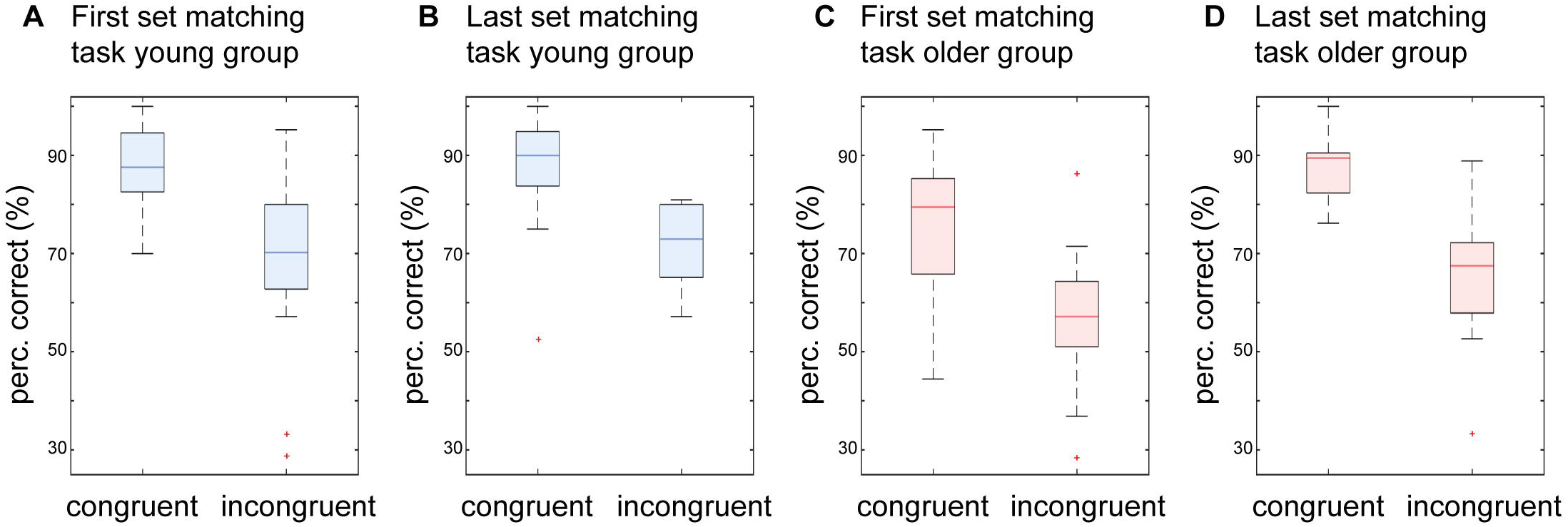
Detection accuracy of congruent vs. incongruent stimulus pairs. Boxplots of detection accuracy for congruent vs. incongruent stimulus pairs for the two groups in the first and the last set of the matching task. **A**: Performance for the first set of the matching task in young participants, **B**: Same as A but for the last set of the matching task in young participants, **C**: Performance for the first set of the matching task in older participants, **D**: Same as C but for the last set of the matching task in older participants.

Adjustment of stimulus intensities increased mean accuracy of pattern detection in the older group (see 3.4). Analyses for congruent vs. incongruent stimulus pairs showed that this effect was driven by better detection of congruent patterns. With increased stimulus intensity, there was a significant increase in the detection of congruent stimulus pairs (*t*(17)=-3.48, *p<*0.01), while detection of incongruent pairs was not significantly enhanced (*t*(17)=-2.04, *p*=0.12). Due to this asymmetric benefit, the congruency effect increased to 23% (88.12% vs. 65.47%, *p*<0.001; see Figure 3D) in the last set of the matching task in the older group.

## Discussion

This study aimed to investigate performance differences in crossmodal integration between young and healthy older adults. The data show that older participants performed worse in unimodal tactile pattern recognition and had higher unimodal detection thresholds. The main finding was that in the crossmodal condition older participants showed higher thresholds compared to the unimodal condition, while young participants showed a stable performance. However, the performance of older participants could be enhanced by further increasing stimulus intensity. This effect was driven by higher detection rates for congruent stimulus pairs, while detection of incongruent pairs was not significantly enhanced. These data support the concept of an increased relevance of congruency effects in older adults in crossmodal tasks.

Confirming our first hypothesis, older participants showed significantly higher thresholds for unimodal tactile and visual pattern recognition than the young. This is in line with previous findings and most likely caused by age-related decline of sensory organs (32,39). The data indicate that healthy older adults are able to perform at a comparable level of accuracy but require higher stimulus intensities (58).

Contrary to our second hypothesis, the data showed that in the older group required stimulus intensities were significantly higher for crossmodal pattern matching compared to the unimodal conditions. Young participants performed significantly better in the crossmodal task at the individually defined perception thresholds. Required stimulus intensities in the young group did not differ between the unimodal and the crossmodal condition. However, even in the complex visuo-tactile matching task older participants were able to reach the same level of performance as in the unimodal detection task. This enhancement of performance with increased stimulus intensities was driven by better detection of congruent stimulus pairs, while detection of incongruent stimuli did not improve, resulting in a numerically stronger congruency effect (23%) than in young adults (18%).

Our data suggest that the previously observed benefit of crossmodal integration in older adults is not necessarily driven by the crossmodal nature of the task but, rather, by congruency of the stimulus materials (21,22). It has been shown that stimulus congruence in crossmodal stimulation can have great impact on behavior compared to unimodal stimulation. The so called ‘congruency effect’ is thought to facilitate cognitive processing and to counteract age-related unimodal shortcomings (39,59,60). This congruency effect has also been shown to apply for crossmodal congruent vs. incongruent information perceived through various modalities (21,36). Our data show congruency effects for congruent vs. incongruent stimulation in young and older participants. At the individual unimodal perception thresholds, congruency effects in the crossmodal task are similar in size in the young and the older group. When performance levels are adjusted, congruency effects drive the improved results in older participants in the crossmodal integration task.

Opposing our initial hypothesis, we did not find evidence for a simple enhancement of crossmodal integration in older participants. There are different reasons that might account for these findings.

Earlier studies indicated that older adults do not benefit from crossmodal stimulation in very complex tasks involving sensory as well as higher-order cognitive processes (15,16,32). Laurienti et al. (2006) found greater performance gains in an older compared to a young group in the crossmodal vs. the unimodal condition. In contrast to our task, they used unequivocal stimulus material, namely colored dots presented on a computer screen and (spoken) color words presented via loud speakers. Poor performance of older participants in our visuo-tactile matching task might be due to increased complexity and task demands of the crossmodal matching compared to unimodal pattern detection. In addition to the complexity of the stimulus material, participants had to pay attention to visual and tactile stimulation concurrently and identify patterns separately in both modalities before comparing them. This might be seen as a worst-case scenario for crossmodal integration. Integration of the two stimuli is not facilitated in a bottom-up manner but has to be actively controlled.

As cognitive top-down-control mechanisms such as attention tend to decline with age and lead to processing difficulties of incoming stimuli (11), this might be another reason for the poor performance of older participants in the crossmodal condition. In line with this, the matching task can also be viewed as a working memory task, as mental representations of patterns have to be compared with each other. The ability to hold information in memory while manipulating it decreases with age (10). Thus, decline of working memory capacity might be another reason for the poor performance of older participants. Taken together, the mechanisms that are thought to lead to enhanced crossmodal integration in older adults, such as the increase of baseline noise (32), general cognitive slowing (29) or inverse effectiveness associated with sensory deficits (15) do not seem to apply in our experimental setting which might be considered a scenario with rather complex task demands.

In summary, older participants performed worse in a complex visuo-tactile matching task at the individual perception thresholds. We do not find behavioral evidence for a simple, compensatory enhancement of crossmodal integration in the older group. However, even in this complex task older participants were able to perform at a comparable level with young adults when higher stimulus intensities were offered. The relative improvement in performance after this adjustment of stimulus intensities was driven by better detection of congruent stimulus pairs. The behavioral benefit of congruent stimulus material in this complex crossmodal scenario underlies the importance of congruency effects.

These findings might have implications for future applications of crossmodal tasks and scenarios. Paying attention to more than one modality and basing one’s decision on a wider range of cues has been suggested to compensate for impaired unisensory processing (25). Destructive consequences of sensory deterioration caused by cognitive and physiological changes are thought to be alleviated this way. Following this idea, Laurienti et al. (2006) suggested the use of crossmodal everyday life gadgets and multisensory training strategies for older adults. The current results add certain limitations to the idea of crossmodal integration as a compensation mechanism for age-related impairments. These limitations include the type and familiarity of stimuli and cognitive demands of a task. Complex cognitive tasks seem to lower the older adults’ capacity to compensate impairments. In this context, the observed benefit of congruent stimulus material might be exploited in future studies and practical applications. To use effects of crossmodal integration in everyday life, congruent information with high stimulus intensities should be delivered through the target modalities. As one of the most important endeavors in this field is to identify means to support older adults to maintain mental health and independent living, congruency effects might be one asset to help older adults master cognitively demanding tasks or to cope with complex scenarios. Congruency effects might also be used to develop strategies for care of disabled older adults as, for example, in neurological rehabilitation.

There are some limitations to the current work. Due to high task demands, there was a high dropout rate in the older group. Therefore, the older participants in our study represented a rather well performing subpopulation. In a more population based older group, the found effects might even be larger. Furthermore, variance in the older participants’ performance was larger compared to the young group. Heterogeneity in older adults is likely to occur with respect to sensory and cognitive impairments. Moreover, highly relevant behavioral and physiological changes not only occur from young to old, but also in higher age (e.g. 7). Considering these aspects, other studies divided participants into young, young-old and old-old. This approach offers the advantage of a more detailed view on the evolution of age-related changes and differences within the older population. Future studies investigating effects of crossmodal integration in older adults might consider recruiting more than two groups.

There are limitations to the current work. In the data presented here, variance in the older participants’ performance was larger compared to the young group. Heterogeneity in older adults is likely to occur with respect to sensory and cognitive impairments. Moreover, highly relevant behavioral and physiological changes occur between the age of 65 and 80 years, merged as one ‘older group’ in the current study (e.g. Poliakoff et al., 2006). Considering these aspects, other studies divided participants into young, young-old and old-old. This approach offers the advantage of a more detailed view on the evolution of age-related changes and differences within the older population. Future studies investigating effects of crossmodal integration in older adults might consider recruiting more than two groups.

## Statement of Ethics

The study was conducted in accordance with the Declaration of Helsinki and was approved by the local ethics committee of the Medical Association of Hamburg (PV5085). All participants gave written informed consent.

## Disclosure Statement

The authors have no conflicts of interest to declare.

## Funding Sources

This work was funded by the German Research Foundation (DFG) and the National Science Foundation of China (NSFC) in project Crossmodal Learning, SFB TRR169/A3/B1/B4 and by the German Research Foundation (DFG) in project SFB 936/A3/C1/Z1.

## Author Contributions

FLH: study design, data acquisition, data analyses, interpretation, preparation of manuscript. CH: data acquisition, data analyses, interpretation, preparation of manuscript. LK: study design, interpretation, revision of manuscript. FG: study design, interpretation, revision of manuscript. AKE: study idea, interpretation, revision of manuscript. FCH: study idea, revision of manuscript GX: study idea, interpretation, revision of manuscript. CG: study idea, study design, interpretation, revision of manuscript.

